# A computational approach for mapping heme biology in the context of hemolytic disorders

**DOI:** 10.1101/804906

**Authors:** Farah Humayun, Daniel Domingo-Fernández, Ajay Abisheck Paul George, Marie-Thérèse Hopp, Benjamin F. Syllwasschy, Milena S. Detzel, Charles Tapley Hoyt, Martin Hofmann-Apitius, Diana Imhof

## Abstract

Heme is an iron ion-containing molecule found within hemoproteins such as hemoglobin and cytochromes that participates in diverse biological processes. While its unlimited supply has been implicated in deleterious processes in several diseases including malaria, sepsis, ischemia-reperfusion, and disseminated intravascular coagulation, little is known about its regulatory and signaling functions. A majority of the computational research to elucidate these functions has been purely data-driven due to the absence of curated pathway resources, which have proven useful in the computational study in other indications. Here, we present two resources aimed to exploit this unexplored information to model heme biology. The first resource is an ontology covering heme-specific terms not yet included in standard controlled vocabularies. Using this ontology, we curated and modeled a corpus of 46 scientific articles to generate a mechanistic knowledge graph representing the heme’s interactome for that particular literature. Finally, we demonstrated the utility of these resources by investigating the role of heme in the Toll-like receptor signaling pathway. Our analysis proposed a series of crosstalk events that could explain the role of heme in activating the TLR4 signaling pathway. In summary, the presented work opens the door for the scientific community to explore in more detail the published knowledge on heme biology.

## 1 Introduction

Heme is an iron ion-coordinating porphyrin derivative essential to aerobic organisms (Zhang, 2011). It plays a crucial role as a prosthetic group in hemoproteins involved in several biological processes such as electron transport, oxygen transfer, and catalysis (Warren & Smith, 2009; Zhang, 2011; Poulos, 2014; Kühl & Imhof, 2014). Besides its indispensable role in hemoproteins, it can act as a damage-associated molecular pattern leading to oxidative injury, inflammation, and, consequently, organ dysfunction (Jeney, 2002; Dutra & Bozza 2014; Wagener et al., 2003; Larsen et al., 2012). Plasma scavengers such as haptoglobin and hemopexin bind hemoglobin and heme, respectively, thus keeping the concentration of labile heme at low concentrations (Smith & McCulloh, 2015). However, at high concentrations of hemoglobin and, consequently heme, the scavenging proteins get saturated, resulting in the accumulation of biologically available heme (Soares & Bozza 2016). With respect to hemolytic diseases, the formation of labile heme at harmful concentrations has been a subject of research for some years now (Roumenina et al., 2016; Soares & Bozza, 2016; Gouveia et al., 2018).

Biomedical literature provides a massive potential source of heterogeneous data that is dispersed through hundreds of journals, making substantial knowledge unseen by the healthcare community and individual researchers. With the introduction of new technologies and experimental techniques, researchers have made significant advances in heme-related research and its role in the pathogenesis of numerous hemolytic diseases such as sepsis (Larsen et al., 2010; Effenberger-Neidnicht & Hartmann 2018), malaria (Ferreira et al., 2008; Dey et al., 2012) and beta-thalassemia (Vinchi et al., 2013; Conran, 2014; Garcia-Santos et al., 2017). The majority of the results are scattered and published as unstructured free-text, or at best, represented in tables and cartoons representing the experimental study or biological processes and pathways. Thus, it is crucial to develop new strategies that capture and exploit this knowledge to better understand the mechanistic role of the heme molecule in hemolytic disorders.

Biological knowledge formalized as a network can be used by clinicians as research and information retrieval tools, by biologists to propose *in vitro* and *in vivo* experiments, and by bioinformaticians to analyze high throughput *-omics* experiments (Catlett et al., 2013; Ali et al., 2019). Further, they can be readily semantically integrated with databases and other systems biology resources to improve their ability to accomplish each of these tasks (Hoyt et al., 2019). However, enabling this semantic integration requires to organize and formalize the knowledge using dedicated controlled vocabularies and ontologies. Although this endeavor involves significant curation efforts, it is key to the success of the subsequent modeling steps. Therefore, in practice, knowledge-based disease modeling approaches have only been conducted for major disorders such as cancer (Kuperstein et al., 2015) or neurodegenerative disorders (Fujita et al., 2013; Mizuno et al., 2012). In summary, while the scarcity of mechanistic information and the necessary amount of curation often hinder launching the aforementioned approaches, modeling and mining literature knowledge provides a holistic picture of the field of interest that can be used for a wide range of applications including hypothesis generation, predictive modeling and drug discovery.

Here, we present two resources aimed to assemble mechanistic knowledge surrounding the metabolism, biological functions, and pathology of heme in the context of selected hemolytic disorders. The first resource is an ontology formalizing heme-specific terms not covered until now by other standard controlled vocabularies. Furthermore, we present a heme knowledge graph (HemeKG), i.e. a network comprising over 700 nodes and over 3,000 interactions as the first attempt to start modelling the knowledge from a collection of more than 20,000 heme-related publications. Finally, we demonstrate both resources by analyzing the crosstalk between heme biology and the TLR4 signaling pathway. The results of this analysis suggest that the activation profile for labile heme as an extracellular signaling molecule through TLR4 is not remarkably distinct from the one established via LPS (lipopolysaccharide) as a signaling molecule and induces cytokines and chemokines, however, the underlying molecular mechanism and individual pathway effectors are not fully understood and need further exploration.

## 2 Materials and Methods

This section describes the methodology used to generate the mechanistic knowledge graph and its supporting ontology. Subsequently, it outlines the approach followed to conduct the pathway crosstalk analysis. A schematic diagram of the methodology is presented in **Figure 1**.

**Figure 1.**
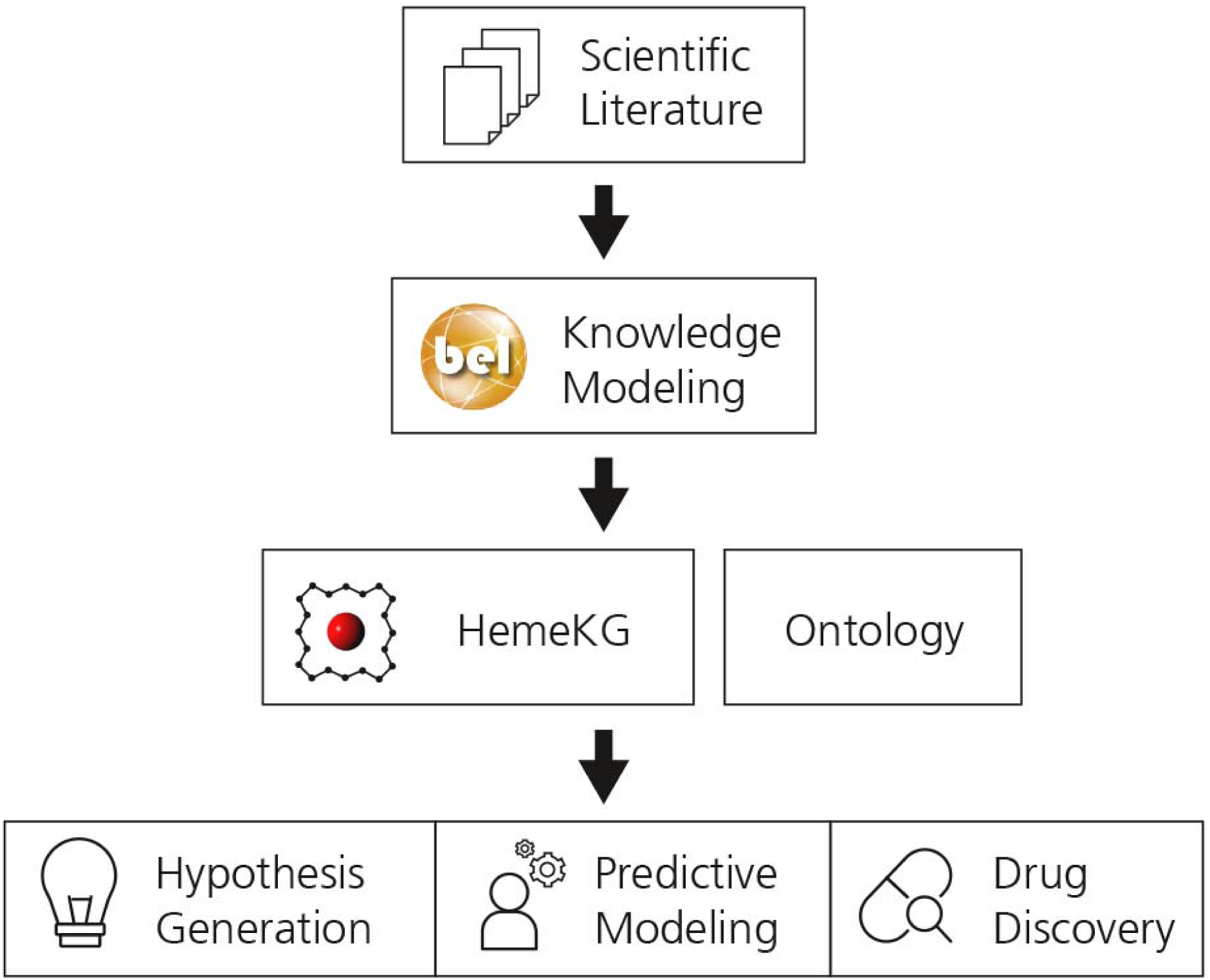
The workflow used to generate the supporting ontology and HemeKG. The first step involves the selection of relevant scientific literature. Next, evidence from this selected corpus are extracted and translated into BEL to generate a computable knowledge assembly model, HemeKG. In parallel to the modeling task, an ontology to support knowledge extraction of articles about the heme molecule was built. Finally, HemeKG can be used for numerous tasks such as hypothesis generation, predictive modeling and drug discovery.

### 2.1 Knowledge modeling

Knowledge was extracted from selected articles (40 original research and 6 review articles) using the official BEL curation guidelines from http://openbel.org/language/version_2.0/bel_specification_version_2.0.html and http://language.bel.bio as well as additional guidelines from https://github.com/pharmacome/curation.

Evidence from the selected corpus was manually translated into BEL statements together with their contextual information (e.g., cell type, tissue and dosage information). For instance, the evidence “Heme/iron-mediated oxidative modification of LDL can cause endothelial cytotoxicity and – at sublethal doses – the expression of stress-response genes” (Nagy et al., 2010) corresponds to the following BEL statement:

~~~
SET Cell = “endothelial cell”
a(CHEBI:”oxidised LDL”) pos bp(MESH:”Cytotoxicity, Immunologic”)
~~~

### 2.2 Generation of a supporting ontology

During curation, an ontology was generated to support the standardization of domain-specific terminology encountered during the curation of articles related to the heme molecule. It comprises terms not present in other controlled vocabularies such as ChEBI (Degtyarenko et al., 2007) for chemicals, or Gene Ontology (GO; (Ashburner et al., 2000)) and Medical Subject Headings (MeSH; (Rogers, 1963)) for pathologies. Each term was checked by two experts in the field assisted by the Ontology Lookup Service (OLS; (Cote et al., 2010)) to avoid duplicates with other terminologies or ontologies. Furthermore, we required that each entry had the following metadata: an identifier, a label, a definition, an example of usage in a sentence, and references to articles in which it was described. Furthermore, a list of synonyms was also curated in a separate file to facilitate the use of the ontology in annotation or text mining tasks. The supporting ontology is included in the **Supplementary Files** and can also be found at https://github.com/hemekg/ontology.

### 2.3 Analyzing pathway crosstalk between heme and the Toll-like receptor signaling pathway

Crosstalk analysis aims to study how two or more pathways communicate or influence each other. While there exist, numerous methodologies designed to investigate pathway crosstalk, the majority of these approaches exclusively quantify such crosstalk based on the overlap between a pair of pathways without delving into the nature of the crosstalk (Donato et al., 2013). In this section, we demonstrate how combining the knowledge from HemeKG with a canonical pathway reveals mechanistic insights on the crosstalk between two different pathways.

Due to the amount of effort required to manually analyze crosstalk across multiple pathways, we conducted a pathway enrichment analysis on three pathway databases (i.e., KEGG, (Kanehisa et al., 2016); Reactome, (Fabregat et al., 2017); WikiPathways, (Slenter et al., 2017) to identify pathways enriched with the gene set extracted from the entire Heme knowledge map. The enrichment analysis evaluated the over-representation of the genes present in HemeKG for each of the pathways in the three aforementioned databases using Fisher’s exact test (Fisher, 1992). Furthermore, Benjamini– Yekutieli method under dependency was applied to correct for multiple testing (Yekutieli and Benjamini, 2001). Manual inspection of the enrichment analysis results revealed that the “Toll-like receptor signaling pathway” was the most enriched pathway in Reactome and WikiPathways, and the third most enriched in KEGG (**Supplementary Table 1**). Therefore, this pathway was selected for study in the subsequent investigation.

First, the three different representations of this pathway were downloaded from each database and converted to BEL using PathMe (Domingo-Fernández et al., 2019). Next, the three BEL networks were combined with the HemeKG network highlighting their overlaps (**Supplementary Figures 1**, in order to specifically analyze these parts of the combined network. Finally, five experts in the field reconstructed the hypothesized pathways from the combined network. The hypothesized pathways were depicted following the guidelines for scientific communication of biological networks outlined by Marai et al. (2019).

## 3 Results

### Building a mechanistic knowledge graph around heme biology in the context of hemolytic disorders

We introduce the first knowledge graph made publicly available to the biomedical and bioinformatics community specifically concerned with heme biology generated using the procedure outlined in the Methods. The presented heme knowledge graph was based on the selection of 40 original research articles and 6 review articles related to heme and its role in several pathways such as Tumor Necrosis Factor (TNF) and nuclear factor kappa-light-chain-enhancer of activated B cells (NF-κB) signaling pathways or the complement and coagulation cascades, through which heme plays a role in hemolysis, inflammation and thrombosis (Dutra and Bozza, 2014; L’Acqua and Hod, 2014; Roumenina et al., 2016; Martins and Knapp, 2018; Vogel and Thein, 2018) (**Figure 2**). The focus of the review articles was chosen due to the relevance of these diseases and complications to large numbers of patients (L’Acqua and Hod, 2014; Litvinov and Weisel, 2016; Roumenina et al., 2016; Effenberger-Neidnicht and Hartmann, 2018). All of these pathologies are known to be interconnected and mapping them in relation to heme is promising for the discovery of yet overlooked links.

**Figure 2.**
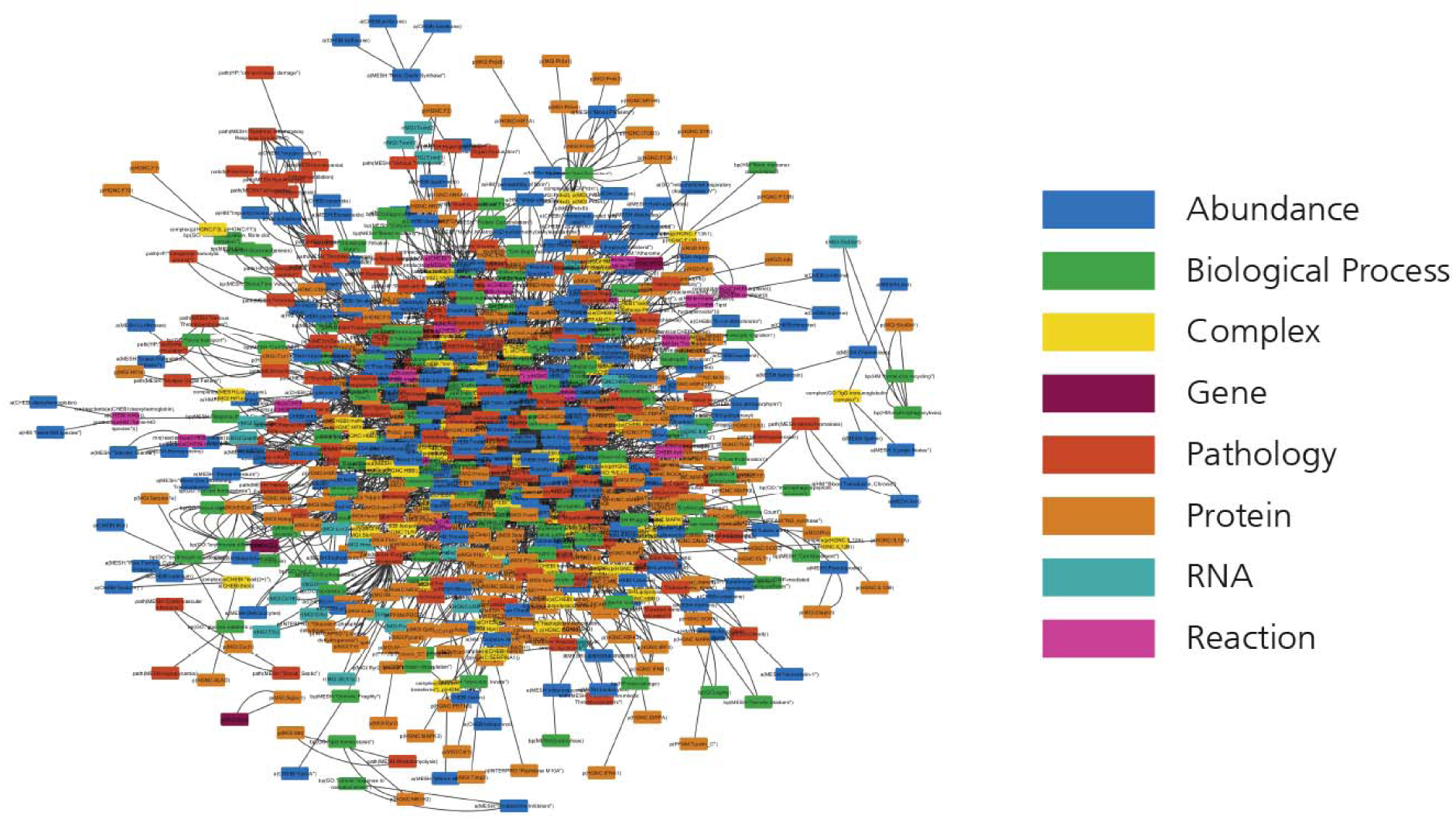
The HemeKG network. Nodes are colored by their different functions in BEL (see legend).

Following the guidelines outlined in the Methods Section, knowledge was extracted and encoded from each of these articles using Biological Expression Language (BEL) due to its ability to represent not only causal, but also correlative and associative relationships found in the literature as well as corresponding provenance and experimental contextual information. The rational enrichment workflow proposed by Hoyt et al. (2019) and Kondratova et al. (2018) that emphasizes curation on low information density nodes in the network was then used to prioritize articles in additional rounds of curation. This strategy was used for four rounds of ten articles to ultimately generate a corpus of 46 curated articles in HemeKG. It contains 775 nodes (**Table 1**) and 3,051 relations (**Table 2**) as well as contextual information ranging from cellular and anatomical localization to different states of the heme molecule (**Supplementary Figure 1**). Annotations such as time point and concentration enabled us to capture time dependencies between entities. By using this contextual information and the multiple biological scales presented in the model, we have not only been able to represent a part of heme’s interactome (**Figure 2**), but also established several links to phenotypes and clinical endpoints. Both represent essential considerations for the design of future clinical studies of hemolytic conditions.

**Table 1.** Summary of unique nodes for each entity class. Each entity class corresponds to the terms formalized in BEL (more information at https://language.bel.bio).

| Abundances | Genes | RNAs | Proteins | Complexes | Reactions | Pathologies | Biological Processes | Total |
| --- | --- | --- | --- | --- | --- | --- | --- | --- |
| 200 | 4 | 25 | 226 | 54 | 17 | 128 | 121 | 775 |

**Table 2.** Summary of relationship classes. Each class corresponds to the relationships formalized in BEL (more information at https://language.bel.bio). The ontological relations class includes the following relationships: has reactant, has product, and has variant.

**Humayun et al., Table 2**
| Increase | Decrease | Positive Correlation | Negative Correlation | Has Component | Association | CausesNo Change | Ontological relations | Total |
| --- | --- | --- | --- | --- | --- | --- | --- | --- |
| 639 | 380 | 1322 | 440 | 113 | 54 | 39 | 64 | 3,051 |

Finally, to facilitate the use of the curated content in this work, BEL documents are bundled with a dedicated Python package that facilitates direct access to the content, provides conversion utilities and allows for network exploration. Both the BEL documents and the Python package are available at https://github.com/hemekg/hemekg.

### Curating a supporting heme ontology

The specificity of our work, together with the lack of contextual terminologies related to heme biology, prompted us to generate a supporting ontology focused on heme. It contains over 50 terms which delineate heme-related biological processes (11), abundances (31), and pathologies (9) that had not yet been included in other standard resources such as Gene Ontology (GO; Ashburner et al., 2000). Building this ontology not only allowed us to describe entities with more expressiveness but also facilitates text mining or annotation tasks related to the heme molecule in the future. The ontology is available at https://github.com/hemekg/ontology.

### Dissection of the crosstalk between heme and TLR using HemeKG

The established heme knowledge graph can be used to study the crosstalk of heme biology in hemolytic disorders with a pathway of interest. In order to select a pathway which highly overlaps with the generated network, we conducted pathway enrichment analysis using three major databases (i.e., KEGG (Kanehisa et al., 2016); Reactome (Fabregat et al., 2017); WikiPathways (Slenter et al., 2017). The results of the enrichment analysis in the three databases pointed to “Toll-like receptor signaling” as the most enriched pathway **(Supplementary Table 1)**. Thus, we proceeded to analyze the crosstalk between this pathway and heme biology by exploring the overlap between HemeKG and the TLR pathways in the three aforementioned databases. Although heme has been linked to numerous Toll-like receptors (TLRs) including TLR2, TLR3, TLR4, TLR7 and TLR9 (Figueiredo et al., 2007; Lin et al., 2010; Dutra and Bozza, 2014; Min et al., 2017), our analysis was prioritized on the most well-documented interaction between heme and TLR4. Heme stimulates TLR4 to activate NF-κB secretion via MyD88 (myeloid differentiation primary response 88)-mediated activation of IKK (see below). Activated IKK promotes the proteolytic degradation of NFKBIA. The phosphorylated IKK complex indirectly activates NF-κB and MAPKs (mitogen-activated protein kinases), such as JNK (C-Jun N-terminal kinase), ERK, and p38 leading to the secretion of tumor necrosis factor alpha (TNF-α), interleukin 6 (IL6), interleukin 1 beta (IL1B), and keratinocyte-derived chemokine (KC) (Dutra and Bozza, 2014). This finally results in the activation of innate immunity and generation of pro-inflammatory factors reflecting the relevance of heme in several disorders, including inflammation and infection.

We first investigated the consensus of the three different representations of the TLR4 signaling pathway (**Figure 3A**). We observed that, overall, all three representations share a high degree of consensus as illustrated in **Figure 3B**. Here, we would like to point out that while KEGG and Reactome present practically alike representations, the WikiPathways representation exhibits slight differences. These differences and complementarities between pathways provide us with more comprehensive views of the studied pathway as illustrated by our previous work (Domingo-Fernández et al., 2019).

**Figure 3.**
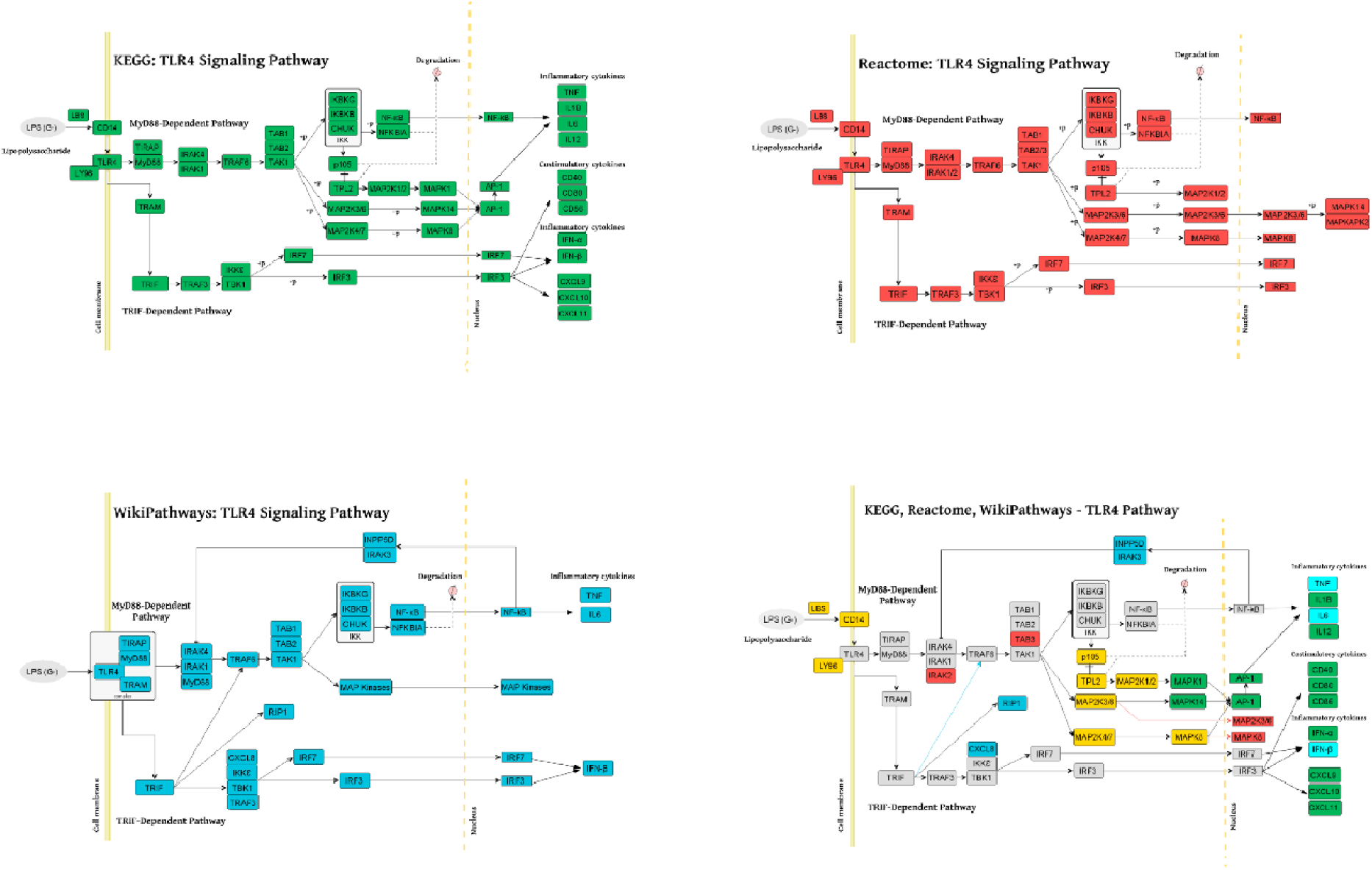
Consensus around the TLR4 signaling pathway in three major pathway databases. A) TLR4 signaling pathway visualization of KEGG, Reactome and WikiPathways. B) Superimposing TLR4 signaling pathway from KEGG, Reactome and WikiPathways. Each color corresponds to the presence of the given node in one or multiple databases (see Legend). MyD88, TAK1, IKK complex, MAP kinases, TNF, NF-κB, TRIF and IRF3 emerged in all three databases and in also in HemeKG. KEGG and Reactome showed identical representations of the TLR4 pathway whereas WikiPathways was different in a way that nuclear NF-κB activates INPP5D-IRAK3 (Inositol Polyphosphate-5-Phosphatase D - Interleukin 1 Receptor Associated Kinase 3) complex which inhibits the activity of IRAK1/IRAK4 (Interleukin 1 Receptor Associated Kinase 1/4).

Secondly, in order to study the overlap between TLR4 signaling pathway and heme biology, we overlaid the consensus network of the pathway with HemeKG (**Figure 4**). Superimposing both networks revealed that MyD88, TAK1, IKK complex, MAP kinases, TNF, NF-κB, TRIF, and IRF3 were present in all three databases as well as in our model. However, several effector molecules, which were found in the three databases, were not found in our heme knowledge graph (HemeKG), e.g., IRAK1, 2, and 4, TRAF6, TAB1-3, and others (**Figure 4A**). Thus, we searched literature reports for specifically these effectors in the context of heme signaling by entering the respective queries in PubMed, as this knowledge might not have been sufficiently covered by the 40 original research articles selected to establish HemeKG.

**Figure 4.**
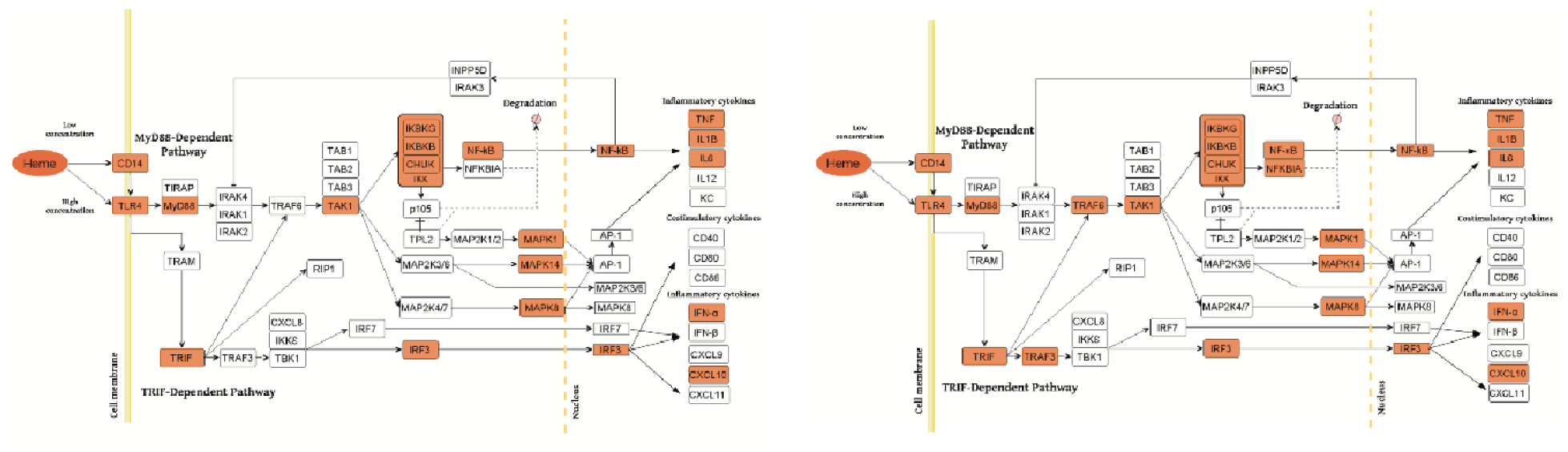
Overlaying the consensus TLR4 signaling pathway in databases with HemeKG (A: Original overlaid network, B: Overlaid network after inclusion of literature evidence for effectors). The orange colored boxes display the common effector molecules between the canonical TLR4 signaling pathway and induced TLR4 signaling pathway stimulated by labile heme. Heme/TLR4 activates the adaptor molecule MyD88. Activated MyD88 promotes the degradation of NFKBIA (NF-κB inhibitor alpha) through phosphorylation of the IKK complex (inhibitor of nuclear factor kappa B kinase complex), thus promoting NF-κB (nuclear factor kappa-light-chain-enhancer of activated B cells) and MAPKs (mitogen-activated protein kinases) stimulation leading to the secretion of TNF-α, IL6, IL1B and KC (keratinocyte-derived chemokine) (Dutra & Bozza, 2014; Fortes et al., 2012). The TRIF (Toll-like receptor adaptor molecule 1) dependent pathway is activated upon signaling of heme through TLR4 leading to the activation of IRF3 (interferon regulatory factor stimulating the secretion of interferons (i.e. IFN-α) and CXCL10 (C-X-C motif chemokine ligand 10) (Dickinson-Copeland et al., 2015). However, the activation profiles for IRAK1/2, TRAF6, TRAM, TRAF3, TBK1/IKK epsilon complex and IRF7 are not yet studied for heme-TLR4 signaling pathway.

The activation profile for labile heme as an extracellular signaling molecule through TLR4 was suggested to be similar to the one established via LPS as signaling molecule from standard pathway databases (Pålsson-Mcdermott and O’Neill, 2004). This pathway begins with the induction of TIRAP (Mal)-associated MyD88 signaling on the one hand (Horng et al., 2002) and TRAM (TICAM-2)-associated TRIF (TICAM-1)-signaling on the other hand (Seya et al., 2005), resulting in the upregulation of pro-inflammatory cytokines and chemokines **(Figure 4**). MyD88 protein as an adaptor has been shown to interact with IRAK (interleukin-1 receptor-associated kinase) proteins 1, 2, and 4 to start the signaling cascade involving TNF receptor associated factor 6 (TRAF6), which is known to activate lkappaB kinase (IKK) in response to pro-inflammatory cytokines. However, in our heme knowledge graph the connections between IRAKs, TRAF6, and TAB proteins were missing (**Figure 4A**). By taking a closer look at these effectors in the context of heme, we found various information for e.g., TRAF6 indicating both a direct and indirect link to heme-induced signaling via TLRs (IJssennagger et al., 2012; Huang et al., 2015; Park et al., 2014; Hama et al., 2012; Meng et al., 2017). In contrast, other effector molecules such as IRAK and TAB proteins (Fig. 4) were not described in heme signaling so far. These findings led us to refine HemeKG in such a way that only those signaling components for which no clear evidence was found still remain white spots on the map (**Figure 4B**).

In addition, the preceding discussion has excluded parameters such as the concentration of labile heme available in the respective environment. This aspect will be particularly important if distinct signaling pathways that may be triggered are dependent on or determined by the concentration of heme. At lower concentrations of heme, TLR4 signaling has been described to be CD14 dependent, whereas at high concentrations of heme TLR4 activation does not require CD14 (Piazza et al., 2010) **(Figure 4)**. Also, there is a need to further investigate whether heme/TLR4 induction of the adapter molecule MyD88 is dependent or independent of TIRAP activation, similarly to the LPS/TLR4 induced TIRAP-associated MyD88 signaling pathway. Furthermore, heme/TLR4 activates a pathway leading to the activation of interferon regulatory factor 3 (IRF3) resulting in the production of interferons e.g., IFN-α (Dutra and Bozza, 2014) and overproduction of C-X-C motif chemokine 10 (CXCL10) (Lin et al., 2012; Dickinson-Copeland et al., 2015). However, the molecular mechanism by which heme/TLR4 induced TRAF3 and, in turn, IRF3/7 activation leads to the secretion of IFN-α and CXCL10 is not yet fully understood adding this pathway to the white spot section of the map (**Figure 4B**). Finally, also the introduction of non-canonical pathways and receptor crosstalk-triggered cascades go beyond the scope of this work, opening opportunities for future studies on heme signaling.

## 4 Discussion

We have presented HemeKG, a first of its kind mechanistic model in the context of heme biology, that provides a first approach to comprehensively summarize heme-related processes by bringing knowledge from disparate literature together. Furthermore, we have demonstrated how combining the knowledge from the heme knowledge graph with information available in pathway databases provides new insights into the network of interactions that regulate heme pathophysiology.

Since HemeKG was curated using standard ontologies, its content can be linked to the majority of public databases. Therefore, enriching the HemeKG network with external data or incorporating its integrated knowledge into other resources is foreseeable. Furthermore, the variety of formats that our resource can be converted to also facilitates its use by other systems biology tools such as Cytoscape (Shannon, 2003) and NDEx (Pratt et al., 2015). In summary, the characteristics of HemeKG not only make this resource suitable for hypothesis generation as presented in our case scenario but also for clinical decision support as previously demonstrated with other systems biology maps (Ostaszewski et al., 2018). For instance, computational mechanistic models are currently being used in combination with artificial intelligence methods for a variety of predictive applications (Khanna et al., 2018; Esteban-Medina et al., 2019; Çubuk et al., 2019). Instead of contextless canonical pathways as until now (i.e., pathways describing normal physiology), HemeKG could be used for predicting drug response and for drug repurposing in numerous related disorders such as malaria and sepsis. Finally, the supporting ontology built during this work could be used for a broad range of applications from data harmonization to natural language processing.

A potential limitation of this study is that it is constrained to a specific literature corpus as we are aware that the presented knowledge graph only captures a part of a much larger interaction network. This tends to be a common challenge when constructing contextualized maps and is further compounded by the difficulty in assessing the coverage of a network. Furthermore, the bias in the scientific community against publishing negative results must also be acknowledged. A clear example is how the hypotheses of our crosstalk analysis could be complemented by this knowledge gap that could reveal new interesting hypotheses. Thus, future updates in HemeKG, as in any work of this kind, will be required to update its content while prioritizing time and effort (Rodriguez-Esteban, 2015). Further, advanced network-based analyses (Yan et al., 2018; Catlett et al., 2013) could be used to rank heme-related pathways in the context of a given *-omics* dataset.

Although numerous interactions between heme and TLRs have been described in the literature (Lin et al., 2010; Min et al., 2017), their downstream effects have not been contextualized (i.e., presented in a coherent/integrated manner like a knowledge model does). The analysis we have presented focusing on the crosstalk between heme biology and the Toll-like receptor signaling pathway has shed some light on how this crosstalk could be related to heme biology. However, other well-known pathways related to heme also exist that could be investigated by conducting similar analyses in the future.

## 4 Data availability

The datasets and scripts of this study can be found at https://github.com/hemekg.

## 6 Author Contributions

DI, MHA and DDF conceived and designed the study. FH curated the data and conducted the main analysis supervised by AAPG, DI and DDF. MTH, BFS, MSD, and AAPG assisted in selecting the corpora and interpreting the results. CTH designed the curation guidelines and implemented the Python package. DDF, FH, CTH, MTH, BFS, MSD, and DI wrote and reviewed the paper.

## 8 Acknowledgments

The authors would like to thank Sarah Mubeen for proofreading the article, and Amelie Wißbrock for useful scientific discussions.

