## Supplementary File for "A computational approach for mapping heme biology in the context of hemolytic disorders"

Supplementary Material

### 1 Supplementary Data

#### 1.1 Articles comprising heme knowledge model

**Review articles (PubMed identifiers):** 30281034, 24904418, 29956069, 25307023, 28458720, 26875449.

**Research articles (PubMed identifiers):** 29929138, 29603246, 29544683, 20378845, 19276082, 30505280, 30324533, 30248094, 29694434, 29610666, 29522519, 29351418, 29212341, 28716864, 28400318, 28314763, 28143953, 28088643, 27798618, 27515135, 27308950, 27125525, 26974230, 26794659, 26675351, 26475040, 26460266, 26368565, 26337933, 26202471, 25411909, 25301065, 25264713, 24667910, 24630724, 24553061, 24489717, 24486321, 24464629, 23215741.

### 2 Supplementary Figures and Tables

#### 2.1 Supplementary Figures

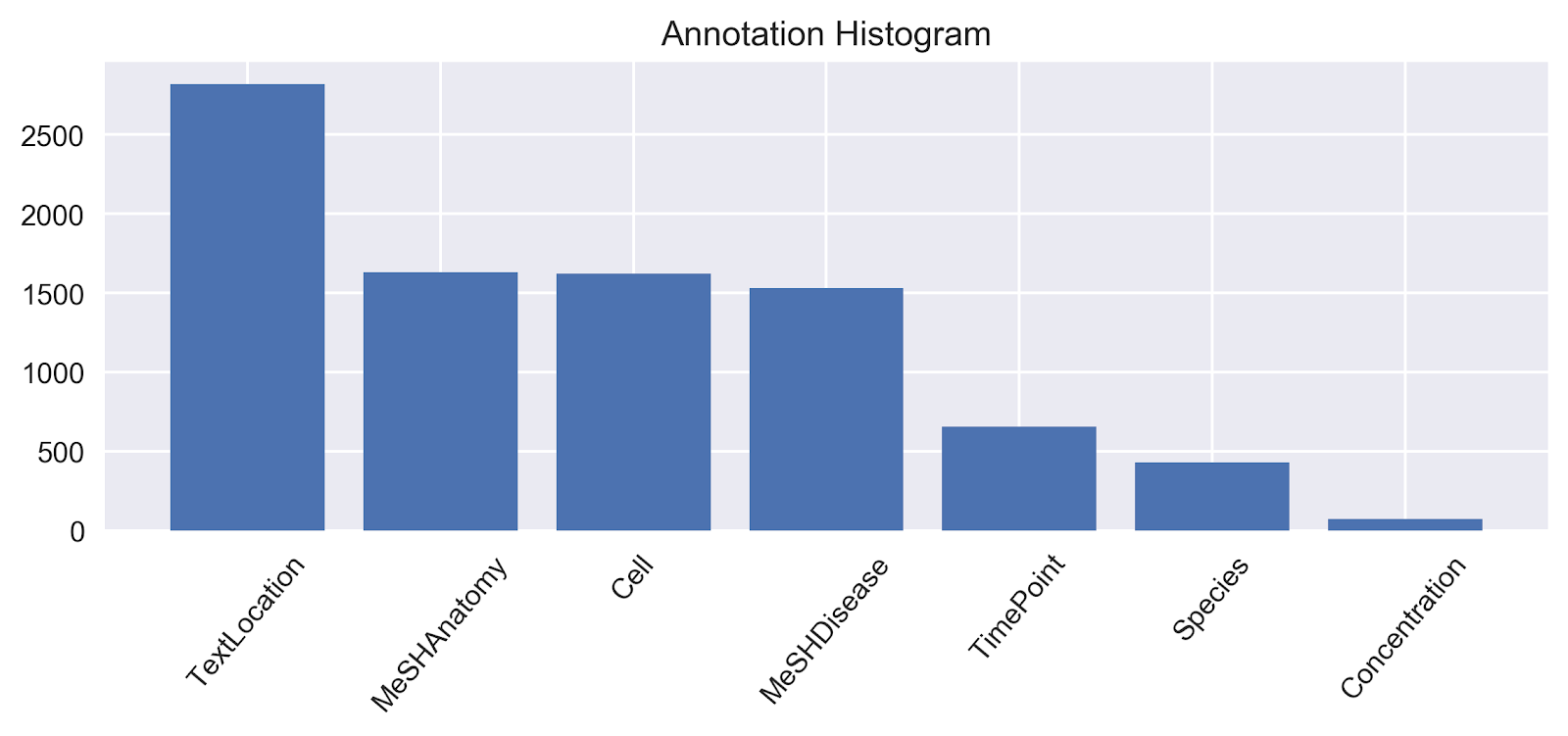

**Supplementary Figure 1.** Annotation statistics. The figure illustrates the statistics for the annotations used while capturing causal relationships from the textual knowledge to generate BEL statements. The textLocation annotation occurs with a frequency of 2,823. Similarly, MeSHAnatomy and MeSHDisease occur with a frequency of 1,635 and 1,533 respectively. Other frequent annotations are cell (1,641), timePoint (660), species (428) and concentration (72).

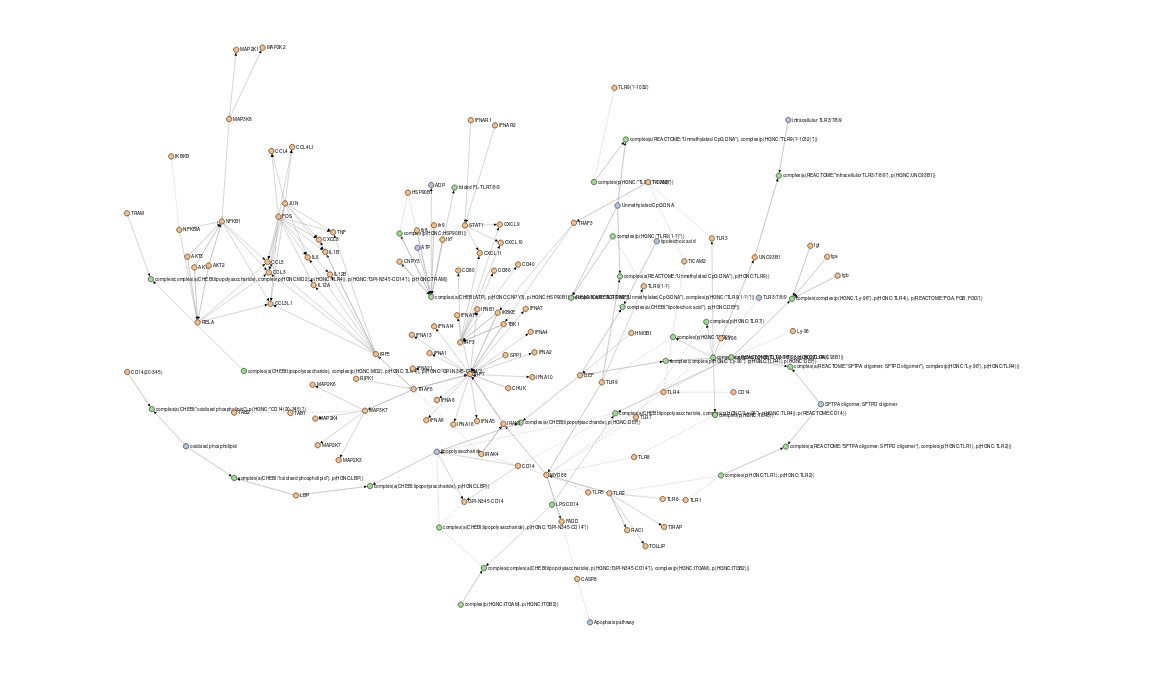

**Supplementary Figure 2.** Toll-like receptor signaling pathways from KEGG, Reactome and WikiPathways in BEL extracted from PathMe [1]. The presented network can be explored and reproduced using the following IPython notebook: <https://nbviewer.jupyter.org/github/hemekg/analysis/blob/master/notebooks/pathways_overlap.ipynb>.

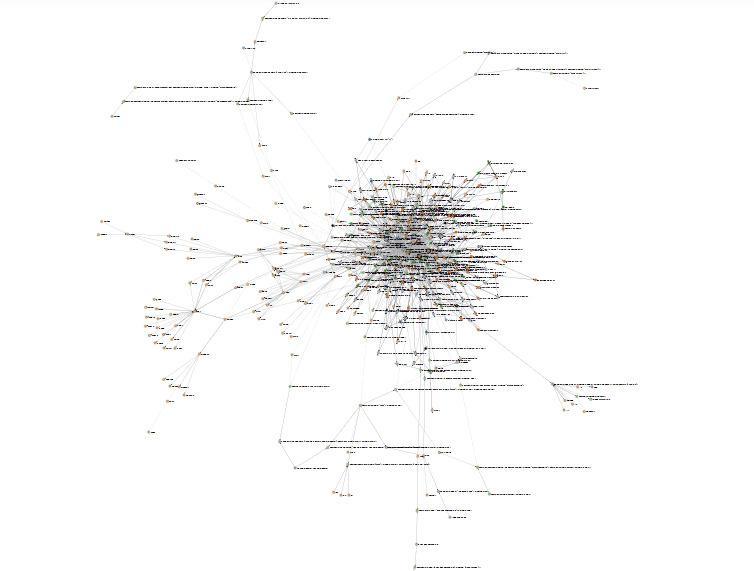

**Supplementary Figure 3.** Overlaying heme knowledge model’s inflammation network with the three representations of the Toll-like receptor signaling pathway from KEGG, Reactome and WikiPathways. The complete overlap to heme knowledge model is not shown due to the large size of the network. The presented network can be explored and reproduced using the following IPython notebook: <https://nbviewer.jupyter.org/github/hemekg/analysis/blob/master/notebooks/pathways_overlap.ipynb>

#### 2.2 Supplementary Tables

| **Pathway** | **Database** | ***p*-value** | ***q*-value** |
| --- | --- | --- | --- |
| Malaria | KEGG | 8.531881e-33 | 2.627819e-30 |
| Complement and coagulation cascades | KEGG | 7.127936e-31 | 1.097702e-28 |
| **Toll-like receptor signaling pathway** | **KEGG** | **1.533560e-30** | **1.574455e-28** |
| Influenza A | KEGG | 5.425161e-28 | 4.177374e-26 |
| Pertussis | KEGG | 6.368401e-27 | 3.922935e-25 |
| **Pathway** | **Database** | ***p*-value** | ***q*-value** |
| **Toll-like Receptor Cascades** | **Reactome** | **2.608813e-25** | **5.598512e-22** |
| Platelet degranulation | Reactome | 4.949028e-23 | 5.310307e-20 |
| Response to elevated platelet cytosolic Ca2+ | Reactome | 9.399391e-23 | 6.723698e-20 |
| Formation of Fibrin Clot (Clotting Cascade) | Reactome | 2.120423e-20 | 1.137607e-17 |
| **Toll Like Receptor 4 (TLR4) Cascade** | **Reactome** | **1.773519e-19** | **7.611943e-17** |
| **Pathway** | **Database** | ***p*-value** | ***q*-value** |
| **Toll-like Receptor Signaling Pathway** | **WikiPathways** | **1.533560e-30** | **7.913169e-28** |
| **Regulation of toll-like receptor signaling pathway** | **WikiPathways** | **1.694942e-29** | **4.372950e-27** |
| Human Complement System | WikiPathways | 6.260545e-29 | 1.076814e-26 |
| Complement and Coagulation Cascades | WikiPathways | 1.133983e-26 | 1.462838e-24 |
| Selenium Micronutrient Network | WikiPathways | 8.509793e-24 | 8.782107e-22 |

**Supplementary Table 1.** Results of the enrichment analysis in the three databases (only top 5 enriched pathways are shown). The enrichment analysis was conducted using ComPath with pathways being downloaded on 1^st^ June, 2019 (Domingo-Fernández *et al*., 2018).

### 3 Supplementary Files

#### 3.1 HemeKG source code

#### <https://github.com/hemekg/hemekg>

#### 3.2 HemeKG raw file

#### <https://raw.githubusercontent.com/hemekg/hemekg/master/hemekg.bel.tsv>

#### 3.3 Ontology and synonyms

### <https://raw.githubusercontent.com/hemekg/ontology/master/terms.csv> <https://raw.githubusercontent.com/hemekg/ontology/master/synonyms.csv>
